# Maternal hypertension and cardiovascular medications dysregulate placental arterial tone

**DOI:** 10.64898/2026.03.24.714086

**Authors:** Teresa Tropea, Elizabeth C. Cottrell, Raianne Wallworth, Nardeen Khalil, Edward D. Johnstone, Jenny E. Myers, Paul Brownbill

**Affiliations:** Maternal & Fetal Health Research Centre, Division of Developmental Biology & Medicine, School of Medical Sciences, Faculty of Biology, Medicine and Health, University of Manchester, Manchester, UK; St Mary’s Hospital, Manchester University Hospital NHS Foundation Trust, Manchester Academic Health Science Centre, Manchester, M13 9WL, UK

## Abstract

**Background:** Antihypertensive and cardioprotective medications are prescribed to pregnant women and include Ca^2+^ channel blockers (CCBs; amlodipine, nifedipine), α- (doxazosin) and β-(labetalol, bisoprolol, nebivolol) adrenergic receptor antagonists, and α-adrenergic receptor agonists (methyldopa). These vasoactive drugs enter the fetal circulation, with unknown effects on the fetoplacental vasculature. We aimed to investigate whether cardiovascular medications modulate human fetoplacental vascular tone, which may impair or enhance placental perfusion.

**Methods:** Chorionic plate arteries (CPAs) were obtained from the placentas of women with normotensive pregnancy (N=28), with unmedicated hypertension (N=14), and those chronically medicated (N=61) with either amlodipine, nifedipine, labetalol or bisoprolol, or a combination of CCBs and labetalol. Using wire myography, *ex vivo* effects of amlodipine, nifedipine, labetalol, methyldopa, doxazosin, bisoprolol and nebivolol were tested in a concentration-dependent manner (10^-11^–10^-5^M) in pre-constricted CPAs isolated from the placentas of normotensive women. Differences in CPA vascular reactivity in response to chronic exposure to hypertension and/or cardiovascular medications was assessed by vasoconstriction to high potassium physiological solution (KPSS; 120mM) and to the thromboxane A_2_ mimetic (U46619; 10^-10^–2×10^-6^M), and relaxation to the nitric oxide donor, sodium nitroprusside (SNP; 10^-10^–10^-5^M).

**Results:** In pre-constricted CPAs isolated from normotensive women, acute exposure to amlodipine, nifedipine, doxazosin and nebivolol promoted significant vasorelaxation (P<0.05). CPAs acutely exposed to labetalol, methyldopa (P<0.05) and bisoprolol (P<0.001) exhibited increased vasoconstriction compared to their respective diluent controls. CPAs from women with chronic hypertension and from those who had chronic labetalol treatment exhibited significantly reduced vasoconstriction to KPSS (P<0.05). CPAs from women with chronic hypertension and exposure to bisoprolol also had significantly attenuated vascular responses to U46619 and SNP (P<0.01 and P<0.01, respectively), compared to normal pregnancy.

**Conclusions:** Maternal hypertension impairs vascular responses of the placenta. Cardiovascular medications prescribed during pregnancy may dysregulate placental vascular function. Further research is warranted to evaluate the relative safety of cardiovascular medications in pregnancy, as their distinct effects on fetoplacental vascular function may have important implications for maternal and fetal outcomes. Mechanistic studies alongside clinical correlations are essential to guide evidence-based prescribing.

## Introduction

Cardiovascular disease is common in pregnant women and is associated with maternal mortality. The spectrum includes cardiac disease and hypertension, both contributing to poor maternal cardiovascular adaptation to pregnancy. Amongst pregnant women who died between 2020-2022, 13% of them were diagnosed with cardiac problems (1).

Pregnancy requires major maternal hemodynamic adaptations to meet fetal metabolic demands, which may challenge the maternal cardiovascular system and contribute to hypertensive disorders (2). Maternal hypertension affects 8-10% of pregnancies worldwide and is diagnosed when blood pressure is greater than 140/90 mmHg (3, 4). Chronic hypertension is defined as occurring before pregnancy or in the first 20 weeks of gestation.

A limited range of antihypertensives, considered safe for the fetus, are prescribed to pregnant women with cardiovascular disease. As the CHIPS and CHAP trials data demonstrated, targeting maternal blood pressure protects the maternal cardiovascular system and prevents the occurrence of severe hypertension (5, 6). Common antihypertensive medications used in pregnancy include labetalol, methyldopa, and calcium channel blockers (CCBs) such as amlodipine and nifedipine (7). Bisoprolol is one of the most common β_1_-adrenergic receptor antagonists prescribed to pregnant women with cardiac disease, including congenital heart disease, cardiomyopathy and arrythmia (8, 9). These medications cross the placental barrier (10) and by entering the fetoplacental circulation, may have detrimental effects on fetal growth and placental function (11).

L-type voltage-gated Ca^2+^ channels regulate vascular tone in the smooth muscle layer. Previous studies showed that L-type VGCCs are functionally expressed in the fetoplacental vasculature and inhibition by CCBs, such as nifedipine, promotes vasorelaxation in chorionic plate arteries (CPAs) (12). Medications targeting the adrenergic system are also recommended to treat maternal hypertension and to minimise cardiac workload in women with cardiac disease.

α- and β-adrenergic receptors mediate the effects of the sympathetic nervous system in response to the catecholamines, noradrenaline and adrenaline, and contribute to the regulation of the cardiovascular system (13). In the systemic circulation, α-adrenergic receptors located on the endothelium, as well as β-adrenergic receptors on both endothelial and smooth muscle cells, have been reported to promote vasorelaxation. In contrast, α-adrenergic receptors expressed on smooth muscle cells mediate vasoconstriction (14). Although the placenta lacks sympathetic innervation, evidence suggests that adrenergic receptors are expressed in the fetoplacental circulation, and that their stimulation and differences in receptor abundance between cell types may evoke different vasoactive effects compared to the systemic circulation (14).

High blood flow and low resistance within the placental vasculature enable adequate delivery of oxygen and nutrients to the growing fetus. Antenatal exposure to mainly β-adrenergic receptor antagonists (15-19) and α_2_-adrenergic receptor agonists (20), has been associated with higher risks of fetal growth restriction (FGR), increased rates of small for gestational age (SGA) newborns and preterm birth. It is unclear whether the increased risks of adverse perinatal outcomes observed in women with cardiovascular disease are attributable to dysregulated placental perfusion associated with the underlying maternal disease or to the medications used to manage these conditions.

Currently, clinicians have minimal guidance to determine which antihypertensive treatment most effectively supports healthy outcomes for mothers and their babies (21). In this study, we tested the hypothesis that administration of antihypertensive and cardiac medications to pregnant women would alter placental vascular function. Using the wire myography system, we tested the acute effects of amlodipine and nifedipine (CCBs); labetalol (α-/β-adrenergic receptor competitive antagonist) methyldopa (α_2_-adrenergic receptor agonist), doxazosin (α_1_-adrenergic receptor antagonist), bisoprolol and nebivolol (β_1_-adrenergic receptor antagonists), on CPAs isolated from the placenta of healthy women. In addition, we assessed the vascular function of CPAs isolated from women with cardiovascular conditions who were either untreated or chronically exposed *in vivo* to either amlodipine, or nifedipine, or labetalol therapies, or labetalol in combination with CCBs, or β1-adrenergic receptor antagonist (bisoprolol) therapy.

## Methods

### Data Availability

The data supporting the findings of this study are available from the corresponding author upon reasonable request. The detailed method, including criteria for the study, can be found in the Supplemental Material.

A total of N=107 study participants was recruited, and the placentas were collected with informed consent following either elective Caesarean section or normal vaginal delivery.

Isolated CPAs were mounted on the wire myography system (22). The study had two distinct objectives. In the first one, arteries obtained from the placenta of normotensive women were pre-constricted with the thromboxane A_2_ mimetic, U46619 (80% maximal effective concentration) and *ex vivo* effects of amlodipine, nifedipine, labetalol, methyldopa, doxazosin, bisoprolol and nebivolol were tested in a concentration-dependent manner (10^-11^–10^-5^M range), based on known (23-29) or estimated (30, 31) fetal venous plasma concentrations (Table S1). In the second objective, vasoconstriction to U46619 (10^-10^–2×10^-6^M), and relaxation (in pre-constricted vessels) to the nitric oxide (NO) donor, sodium nitroprusside (SNP; 10^-10^–10^-5^M), was tested on CPAs isolated from placentas of women with hypertension who either remained untreated throughout their pregnancies or were treated with cardiovascular medications to assess the effect of chronic exposure to hypertension and/or cardiovascular medications on fetoplacental vascular reactivity.

### Statistical analysis

Concentration-responses to SNP and medications are expressed as percentage level of vasoconstriction remaining from the pre-constriction achieved with the U46619 EC_80_ dose. All data are expressed as mean ± SEM and differences between groups analysed by 1- or 2-way ANOVA or as otherwise stated. Statistical significance was defined as P < 0.05 (for details see Supplemental Material).

## Results

Table 1 summarises the clinical characteristics of pregnant women and their pregnancy outcomes (for demographics of pregnant women see Tables S2 Supplemental Material), grouped by maternal disease and/or medication exposure

**Table 1.**
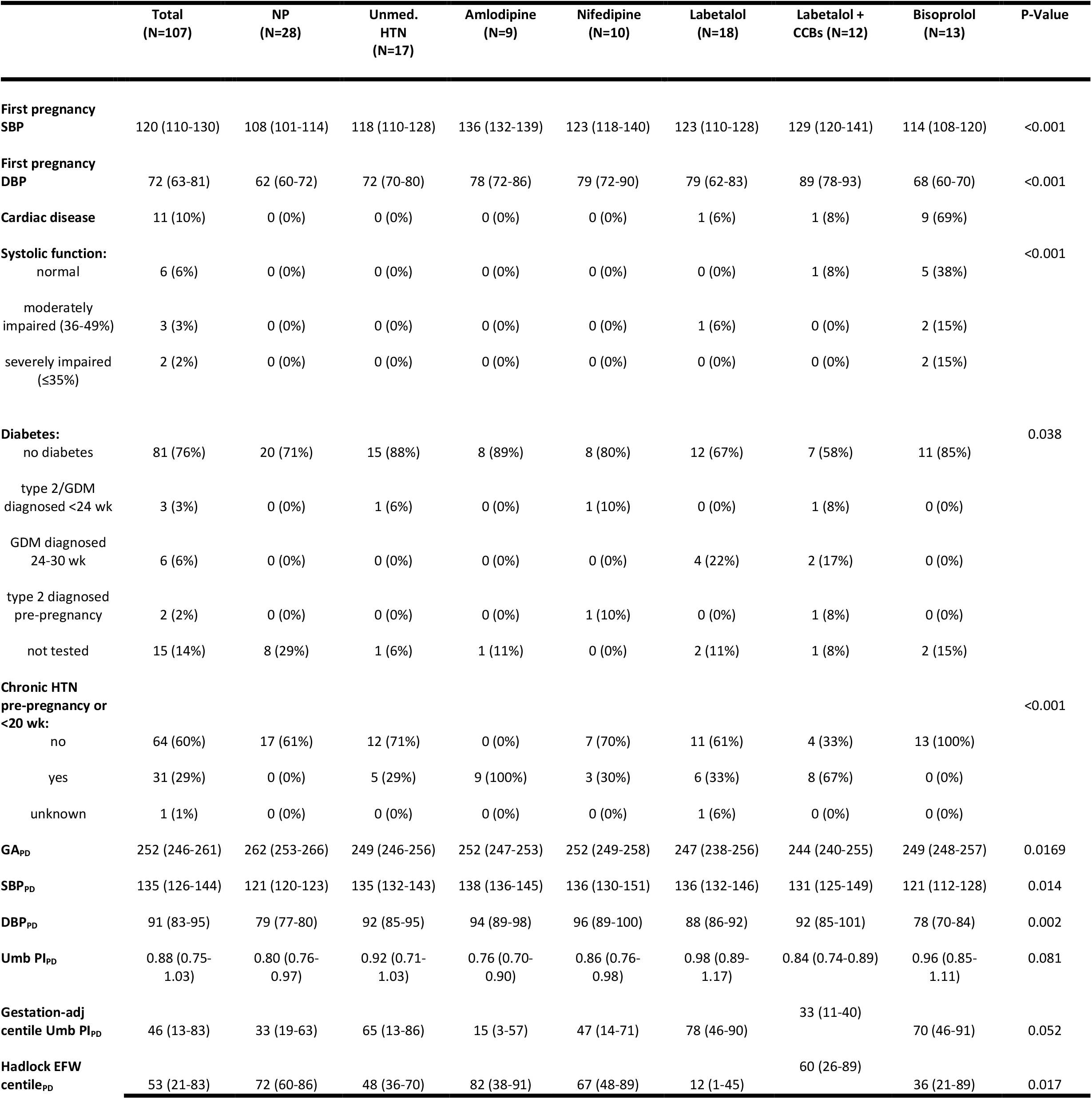
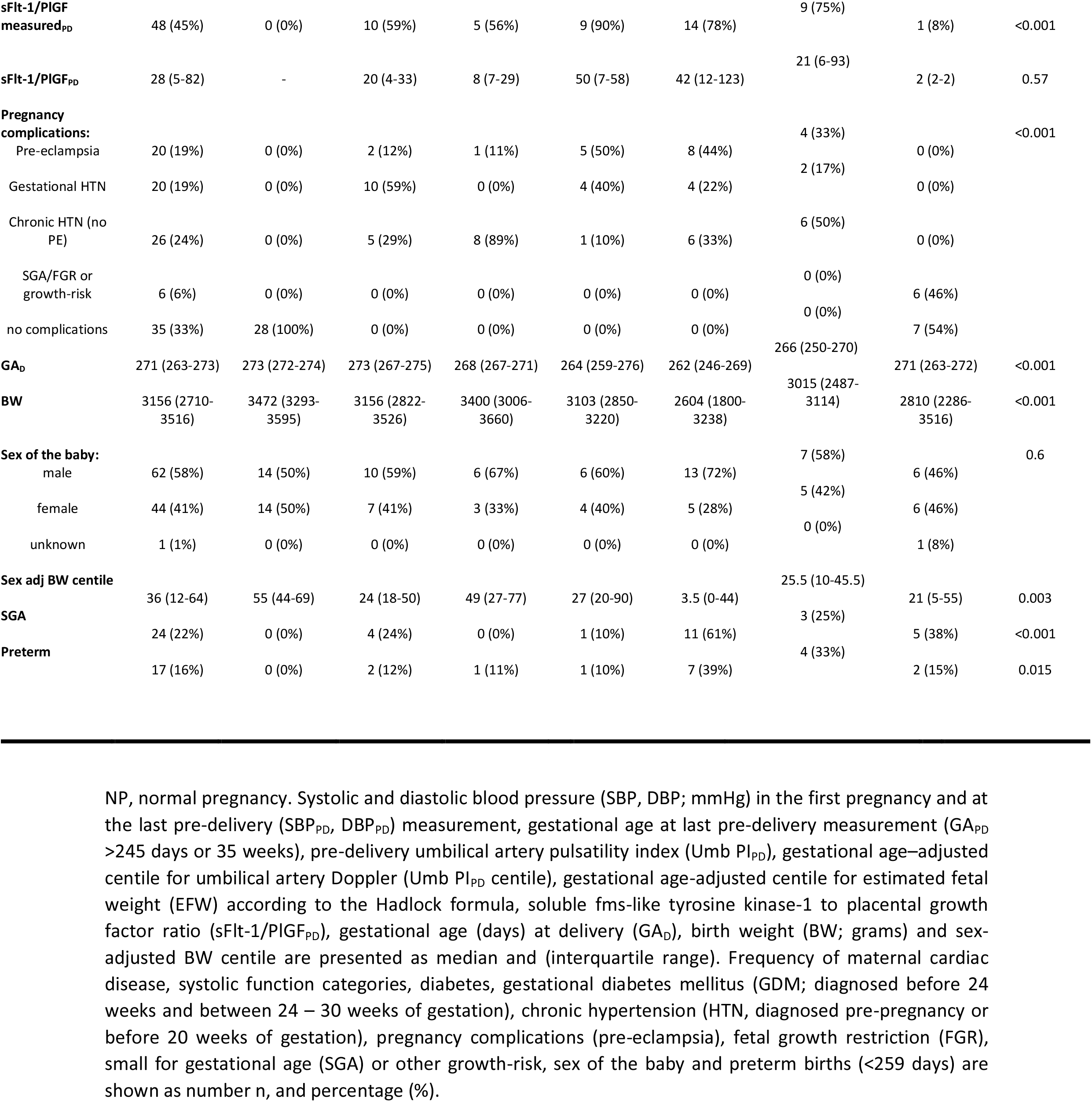
Clinical characteristics and pregnancy outcomes of study participants.

In the total cohort, first pregnancy and pre-delivery blood pressures were higher in the unmedicated and in the medicated hypertensive groups compared to those with no hypertension. 10% of women had cardiac disease, with the highest rate in the bisoprolol group (69%); in the same group, systolic function was moderately and severely impaired in a total of 30% of women. 3% of women had a diagnosis of type 2 and gestational diabetes mellitus (GDM) before 24 weeks, 6% were diagnosed with GDM between 24-30 weeks of pregnancy, and 2% had type 2 diabetes pre-pregnancy. 29% of women had chronic hypertension, with the highest rate in the amlodipine- (100%) and in the labetalol + CCBs-medicated (67%) groups. Pregnancy complications affected 19% of pregnancies in the form of pre-eclampsia (PE), mainly in the nifedipine- (50%) and in the labetalol-medicated (44%) groups.

Umbilical Doppler assessments showed a trend toward higher PI, with median gestation-adjusted umbilical artery PI centiles of 78 (46–90) and 70 (46–91) in the labetalol and bisoprolol groups, respectively. These groups also had the highest rates of SGA (61% and 38%, respectively).

The bisoprolol cohort had the longest exposure time during pregnancy amongst all the medicated groups (Table S3).

Labetalol, whether administered alone or in combination with CCBs, was associated with higher number of preterm births (39% and 33%, respectively).

### Acute *ex vivo* exposure to antihypertensives differentially alters fetoplacental vascular tone

When tested *ex vivo*, the CCBs amlodipine (Figure 1A) and nifedipine (Figure S1), the α1-adrenergic receptor antagonist doxazosin (Figure 1D), and the third generation β1-adrenergic receptor antagonist nebivolol (Figure 1E), promoted significant vasorelaxation in submaximally pre-constricted CPAs isolated from the placenta of normotensive pregnant women. Compared to controls, amlodipine caused 31.9 ± 4.3% maximum vasodilatory effect (Emax; P<0.0001 overall effect vs control), followed by nebivolol (P=0.036 overall effect vs control) 17.9 ± 5.6% Emax, and doxazosin (P=0.0183 overall effect vs control) 14.8 ± 4.6% Emax.

**Figure 1.**
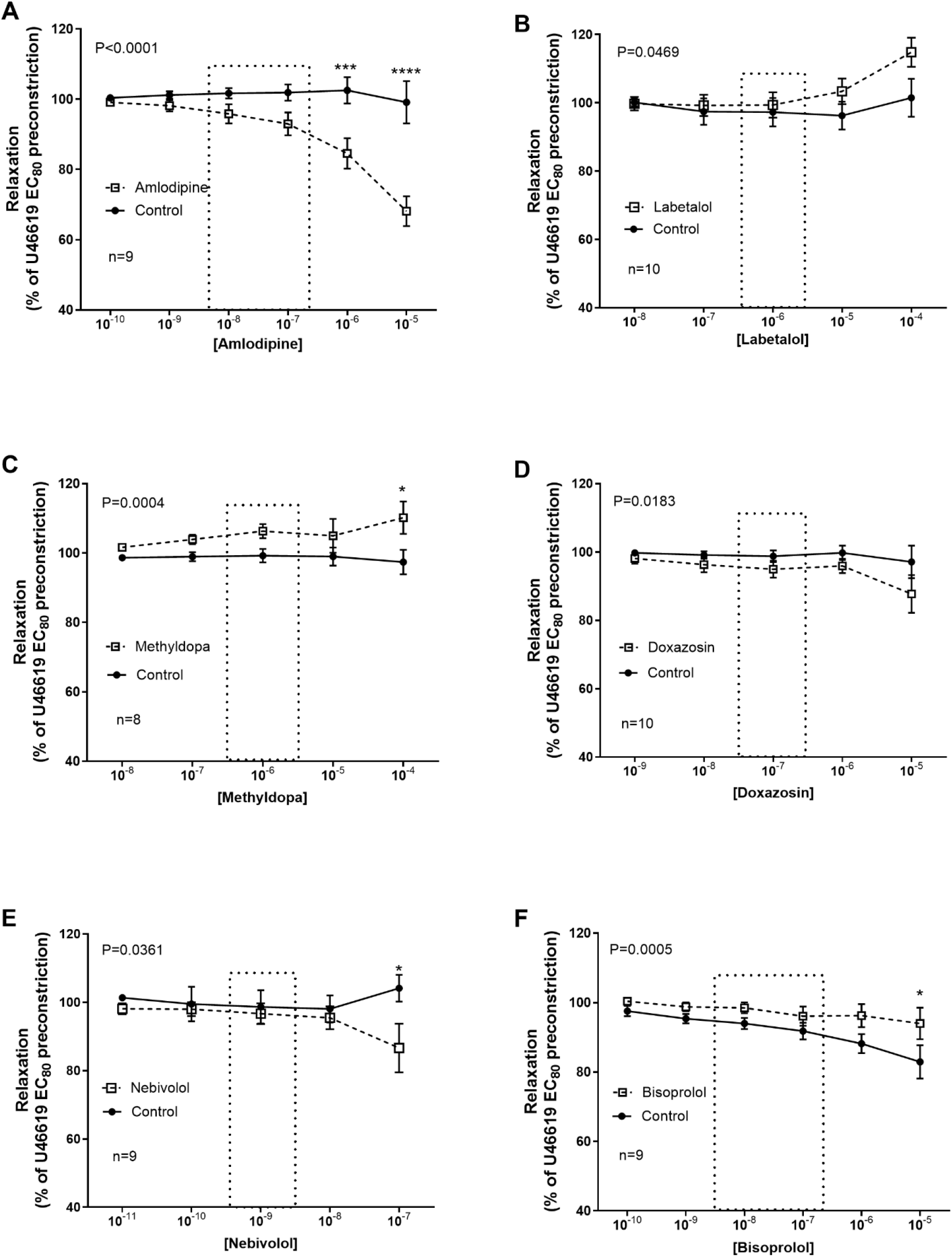
Contrasting effects of different antihypertensive and cardioprotective medications on chorionic plate arteries *ex vivo*. Cumulative concentration-response curves for (A) amlodipine, (B) labetalol, (C) methyldopa, (D) doxazosin, (E) nebivolol and (F) bisoprolol vs their relative diluent controls on pre-constricted fetoplacental arteries. Concentrations shown within the dotted columns represent the predicted in vivo dose exposure. Curves are compared by 2-way ANOVA; P values are the overall effect vs control. N=8-10 per group. *P <0.05, ***P <0.001, ****P <0.0001; Šídák’s multiple comparison test.

The α/β-adrenergic receptor competitive antagonist labetalol (Figure 1B), the α_2_-adrenergic receptor agonist methyldopa (Figure 1C) and the second generation β1-adrenergic receptor antagonist bisoprolol (Figure 1F) exhibited increased vasoconstriction compared to their relative controls, with labetalol (P<0.0469 overall effect vs control) reaching 15.5 ± 4.0% Emax, and methyldopa (P=0.0004 overall effect vs control) causing 13.0 ± 3.0% Emax. Bisoprolol maintained higher level of pre-constriction over time compared to its relative control group (94.0 ± 4.6% vs 82.9 ± 4.8% at the maximum concentration tested; P=0.0005 overall effect vs control).

### Maternal hypertension, but not Ca^2+^ channel blocker therapy, reduces constriction and relaxation responses in *ex vivo* chorionic plate arteries

Compared to the normotensive group (NP), CPAs in the untreated hypertensive group (HTN) had a significantly reduced response to 120 mmol/L K^+^ solution (KPSS vasoconstriction in kPa: NP, 9.2 ± 0.6; HTN 6.8 ± 0.6; P< 0.05 Figure 2A&D). In comparison with the NP group, CPAs from women treated with CCBs did not have significantly altered KPSS responses (amlodipine, 7.5 ± 0.7; nifedipine, 8.0 ± 1.1). Cumulative concentration-dependent vasoconstriction to U46619 was significantly reduced in the HTN and in the CCB groups (U46619 Emax in kPa: NP, 14.8 ± 1; HTN, 10.0 ± 0.8; amlodipine, 10.9 ± 1.1; nifedipine, 11.5 ± 1.4. P<0.0001 Figure 2B and 2E). Vasorelaxation was slightly reduced at SNP concentrations of 10^-7^ and 10^-6^ M in the CCB groups, compared to the NP group, but was not different to the underlying HTN effect (Figure 2C and 2F). Overall, HTN reduced responses to SNP compared with NP (SNP Emax in %: NP, 73.1 ± 4.6; HTN, 62.8 ± 4.3; amlodipine, 68.5 ± 5.69; nifedipine, 62.3 ± 5.7. P = 0.005 Figure 2C and P=0.0006 2F). There was no difference between HTN and CCB groups.

**Figure 2.**
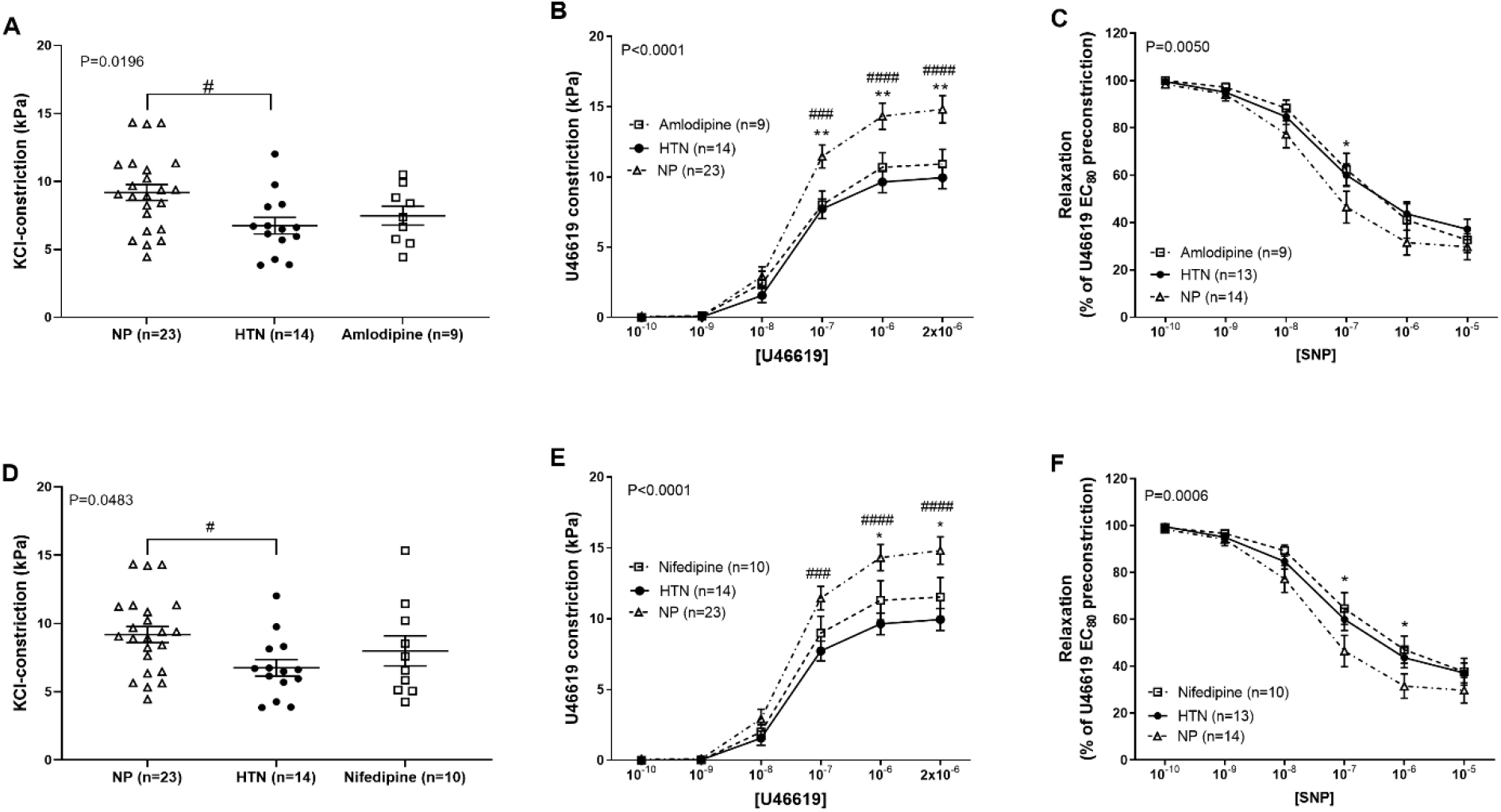
Effects of hypertension and Ca^2+^ channel blocker exposure in women with hypertension on chorionic plate artery vascular responses. Vasoconstriction in response to 120 mM KCl (A, D), and U46619 (B, E); vasorelaxation in response to sodium nitroprusside (SNP; C, F), in chorionic plate arteries isolated from the placenta of women with normal pregnancy (NP), unmedicated hypertension (HTN) and medicated with either amlodipine (A, B, C) or nifedipine (D, E, F). Groups are compared by 1-way ANOVA (A, D) or 2-way ANOVA (B, C, E, F); P values on the graphs are the overall effect. #P < 0.05, ###P < 0.001, ####P < 0.0001, HTN vs NP; *P < 0.05, **P < 0.01, amlodipine or nifedipine vs NP; Tukey’s multiple comparison test.

### Labetalol treatment does not worsen hypertension-associated vascular dysfunction in *ex vivo* fetoplacental arteries

In women with HTN exposed to labetalol during pregnancy, vasoconstriction of CPAs in response to 120 mmol/L K^+^ solution and to U46619 was overall significantly lower compared to the NP group (KPSS vasoconstriction in kPa: 5.9 ± 0.5 P= 0.0002 Figure 3A; U46619 Emax in kPa: 11.1 ± 1.0. P<0.0001 Figure 3B). Compared to the NP group, vasorelaxation to SNP was significantly reduced in the labetalol group (SNP Emax in %: labetalol 68.1 ± 3.4, P=0.0042 Figure 3C). However, there was no significant difference between HNT and labetalol groups.

**Figure 3.**
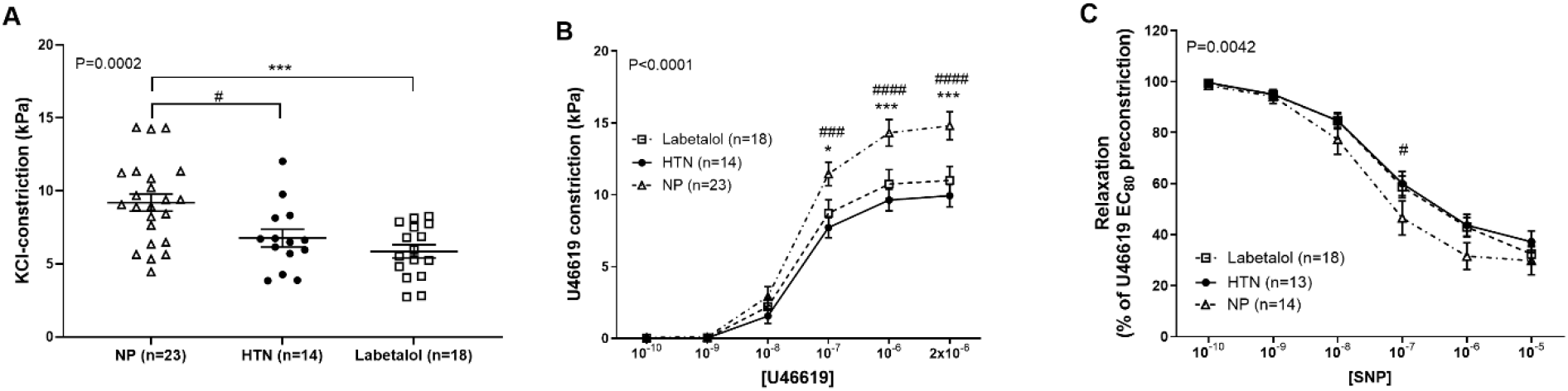
Effect of labetalol exposure in women with hypertension on chorionic plate arteries. Vasoconstriction in response to (A) 120 mM KCl, and (B) U46619; (C) vasorelaxation in response to sodium nitroprusside (SNP), in chorionic plate arteries isolated from the placenta of women with normal pregnancy (NP), hypertension unmedicated (HTN), and medicated with labetalol. Groups are compared by 1-way ANOVA (A) or 2-way ANOVA (B, C); P values on the graphs are the overall effect. #P < 0.05, ###P < 0.001, #### P < 0.0001, HTN vs NP; *P < 0.05, ***P < 0.001, labetalol vs NP; Tukey’s multiple comparison test.

### *In vivo* Ca^2+^ channel blockers combined with labetalol does not further dysregulate the underlying hypertension effect in *ex vivo* fetoplacental arteries

In CPAs isolated from hypertensive women taking a combination of labetalol and CCBs during pregnancy, vasoconstriction in response to 120 mmol/L K^+^ solution was reduced compared to the NP group (KPSS vasoconstriction in kPa: labetalol + CCBs, 8.1 ± 0.8. P= 0.0336 Figure 4A). However, there was no difference between the HTN and labetalol + CCBs-treated group. A significant attenuation of vasoconstriction in response to U46619 (U46619 Emax in kPa: 11.2 ± 1.0. P<0.0001 Figure 4B), and of vasorelaxation to SNP (SNP Emax in %: 68.1 ± 3.3. P=0.0011 Figure 4C) was shown in the labetalol + CCBs group compared to NP. At a concentration of SNP 10^-7^M, comparison between groups showed significant effect of reduced vasorelaxation in both the HTN and the labetalol + CCBs-treated group (Figure 4C).

**Figure 4.**
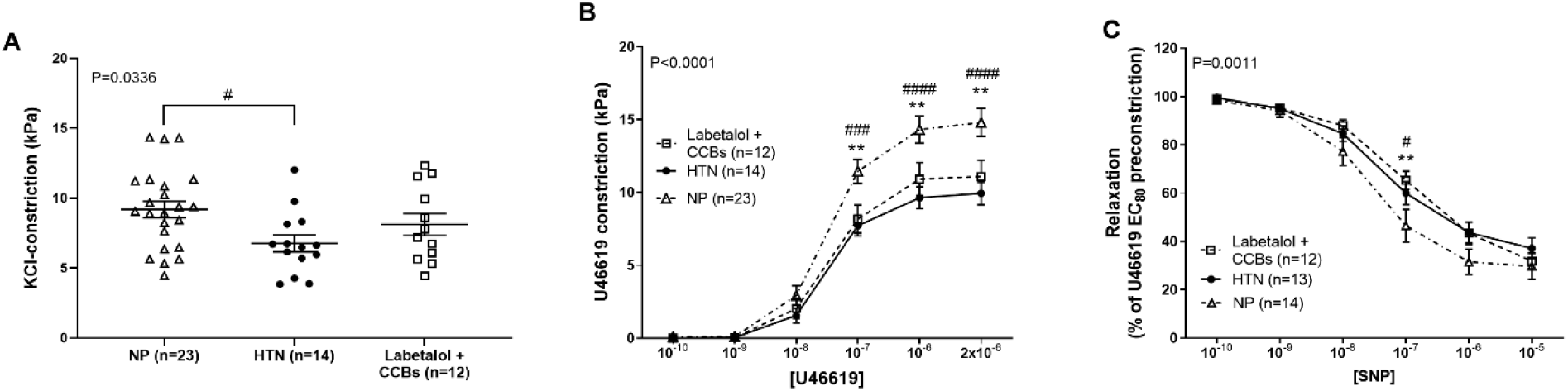
Effect of labetalol in combination with Ca^2+^ channel blocker therapy exposure in women with hypertension on chorionic plate arteries. Vasoconstriction in response to (A) 120 mM KCl, and (B) U46619; (C) vasorelaxation in response to sodium nitroprusside (SNP), in chorionic plate arteries isolated from the placenta of women with normal pregnancy (NP), unmedicated hypertension (HTN), and hypertension medicated with labetalol and Ca^2+^ channel blockers (CCBs, amlodipine, nifedipine). Groups are compared by 1-way ANOVA (A) or 2-way ANOVA (B, C); P values on the graphs are the overall effect. #P < 0.05, ###P < 0.001, ####P < 0.0001, HTN vs NP; **P < 0.01, labetalol + CCBs vs NP; Tukey’s multiple comparison tests.

### *In vivo* bisoprolol exposure diminishes vasoconstriction and vasorelaxation responses in *ex vivo* fetoplacental arteries

Women with cardiac dysfunction included in this cohort were not hypertensive, therefore, this group was compared with CPAs from women with a normotensive pregnancy. In women with cardiac dysfunction, treatment with bisoprolol during pregnancy did not affect vasoconstriction of CPAs in response to 120 mmol/L K^+^ solution (KPSS vasoconstriction in kPa: bisoprolol, 7.7 ± 1.0. P=0.1847 Figure 5A); whereas CPAs from this group exhibited significantly reduced vasoconstriction to U46619 (U46619 Emax in kPa: 12.1 ± 1.3. P=0.0087 Figure 5B) compared to NP. Comparison between groups also showed that chronic bisoprolol exposure significantly reduced vasorelaxation to SNP (SNP Emax in %: 54.3 ± 5.1. P=0.0002 Figure 5C).

**Figure 5.**
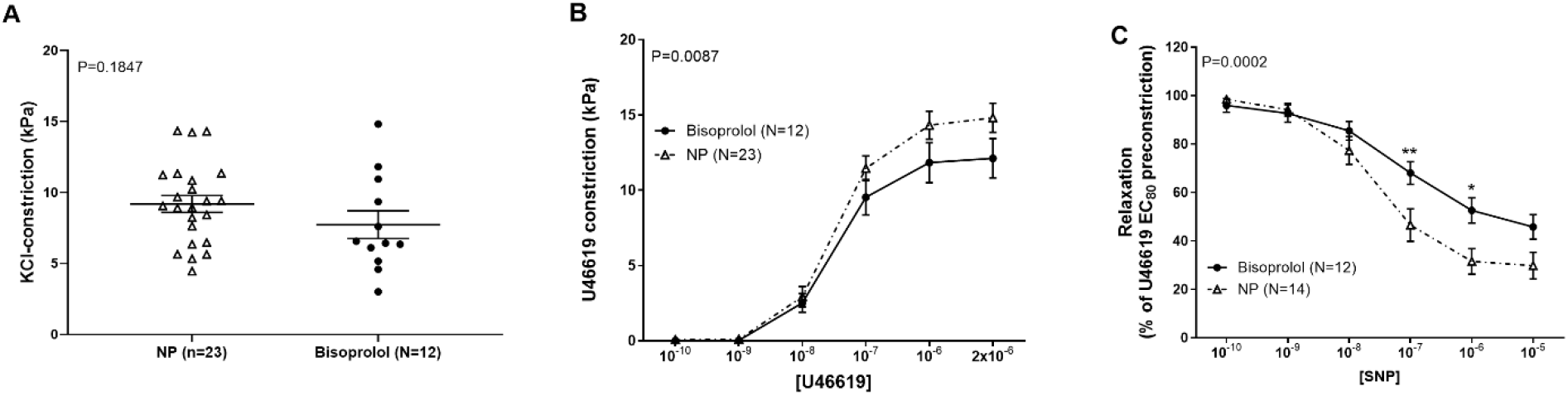
Effect of bisoprolol exposure in women with cardiac dysfunction on chorionic plate arteries. Vasoconstriction in response to (A) 120 mM KCl, and (B) U46619; (C) vasorelaxation in response to sodium nitroprusside (SNP), in chorionic plate arteries isolated from the placenta of women with normal pregnancy (NP) and those with cardiac dysfunction medicated with bisoprolol. Groups are compared by unpaired t-test- (A) or 2-way ANOVA (B, C); P values on the graphs are the overall effect. *P<0.05, **P < 0.01, bisoprolol vs NP; Šídák’s multiple comparisons test.

There was insufficient data to compare the effects of *in vivo* nebivolol exposure on fetoplacental vascular responses. Exclusion of CPA samples obtained from PE participants did not affect data analysis (Figure S2-S4).

## Discussion

To our knowledge, this is the first study to compare the effects of maternal hypertension, and exposure to antihypertensive and cardioprotective medications commonly prescribed in pregnancy, on human fetoplacental vascular function.

Our *ex vivo* data revealed that acute exposure to CCBs promoted vasorelaxation, whereas medications modulating the adrenergic system elicited contrasting effects on CPAs from normotensive healthy pregnancies. Acute labetalol, methyldopa and bisoprolol exposure induced increased vasoconstriction compared to respective diluent controls. In contrast, doxazosin and nebivolol induced significant vasorelaxation. Using CPAs from pregnancies complicated by hypertension and cardiac disease, we found that maternal hypertension *per se* reduced CPA reactivity in response to both high potassium solution and to a thromboxane-mimetic vasoconstrictor. Vasorelaxation to exogenous NO was also attenuated in CPAs from women with hypertensive pregnancy. We observed no major differences between CPAs from the untreated hypertensive group and those treated with different medication classes during pregnancy.

In CPAs from normal pregnancies, L-type CCBs amlodipine and nifedipine induced marked vasorelaxation. Our findings are consistent with previous studies on fetoplacental arteries in our group (12), and in those of others conducted in human internal mammary arteries at a comparable concentration-range (32). In contrast to CCBs, medications targeting the adrenergic system, produced more disparate effects. Compelling evidence showed that the human fetoplacental vasculature expresses adrenergic receptors, and their vascular effects vary depending on the activation/deactivation of specific receptor subtype(s) (14). Exposure to β-adrenergic receptor antagonists, including labetalol, reduces fetoplacental blood flow and increases placental vascular resistance in hypertensive pregnant ewes (33). Maternal labetalol administration increases umbilical artery pulsatility index (PI) in human pregnancies, an effect attributed to increased placental vasoconstriction (34), and supported by evidence from *ex vivo* studies using human placentas perfused with labetalol (35). Whilst nifedipine increases placental perfusion *ex vivo* (36), in humans (37) and in animal models (38), in comparison with the oral treatment with labetalol, there was no significant difference in maternal and neonatal outcomes (39). In contrast to our findings of labetalol and methyldopa, which further increased vasoconstriction in arteries pre-constricted with U46619, previous work has shown vasorelaxing properties of labetalol in human forearm radial arteries rings, partly mediated via activation of calcium-activated potassium channels when tested acutely *ex vivo* (10^-8^ - 10^-4^ M range)(40). Labetalol is a non-selective and competitive β-blocker, with an approximate β- : α-blockade ratio of 7:1 (41). Given the competitive antagonism of labetalol favouring β-blockade, it may produce a stronger inhibitory effect on the endothelial-dependent vasodilatory actions mediated by β-adrenergic receptor activation. The resulting imbalance in the blockade effect, and the higher expression of α-adrenergic receptors in human term fetoplacental arteries (42), may promote α-mediated vasoconstriction, thereby potentiating the initial level of pre-constriction in response to the thromboxane mimetic in the smooth muscle cells. This observation is consistent with previous findings from *ex vivo* dual perfusion studies of the human placenta, where labetalol administration increased the perfusion pressure induced by endothelin-1 and U46619 (35). The α-adrenergic agonism of methyldopa might have contributed to the observed comparable response in the fetoplacental vasculature.

Bisoprolol did not evoke overt vasoconstriction *ex vivo*, but vessels exhibited a sustained level of tension over time relative to the diluent control. Compared to labetalol, bisoprolol is a cardioselective β-blocker with ∼100 times higher affinity for β1-over β2-adrenergic receptors, and with no affinity for α-adrenergic receptors (43). The absence of α-adrenergic agonism might have slightly attenuated the initial pre-constriction, while the β-blocking properties likely prevented vasorelaxation.

In contrast, the α-adrenergic receptor antagonism of doxazosin (44), and the vasodilatory properties of nebivolol (45, 46) promoted relaxation of the fetoplacental vasculature *ex vivo*. Studies have demonstrated the vasodilatory effect of nebivolol in human penile arteries via NO-dependent mechanisms (47) and partly mediated through calcium activated potassium channels in human radial arteries (40). In coronary resistance microarteries of humans and rodents, nebivolol has been shown to target the endothelial β3-adrenergic receptors (46), and to release NO via β2-adrenergic receptors in mouse thoracic aorta (48). Our study was underpowered to investigate the effect of *in vivo* exposure to nebivolol. However, consistent with our findings of acute exposure causing vasorelaxation, human umbilical vein endothelial cells (HUVECs) exposed to plasma from preeclamptic women showed increased NO production in the presence of nebivolol, an effect that was abolished by NO-synthase and β3-adrenergic receptor inhibitors (49).

While endothelial dysfunction and increased production of vasoconstrictors are key factors in the pathophysiology of hypertension (50), vascular smooth muscle cells are the functional effectors of vascular tone modulation (51). In our study, we challenged mechanisms of vasoconstriction and relaxation both converging on vascular smooth muscle cell function. We showed that maternal hypertension was associated with impaired vasoconstriction and reduced endothelium-independent vasorelaxation in CPAs, independent of medication exposure during pregnancy. This suggests that the fetoplacental circulation is altered in pregnancies complicated by maternal hypertension, and that vascular responsiveness to vasoactive agents is different to the systemic circulation potentially due to lower level of sympathetic tone regulation and imbalance between α- and β-adrenergic receptor mediated effects in the fetoplacental vascular bed. Pre-eclampsia developed in 19% of the participants; however, no cases occurred in either the normal or the bisoprolol group. Exclusion of pre-eclampsia cases from our groups did not affect data analysis.

Although current evidence indicates an increased risk of infants being born small for gestational age (SGA) in women with hypertensive disorders of pregnancy, it is uncertain whether this association is exacerbated by antihypertensive medication (52). The Giant PANDA trial, a pragmatic, multicenter, randomized controlled study involving 2,200 women with pregnancy hypertension, aims to generate high-quality evidence to guide the selection of first-line antihypertensive therapy by balancing effective maternal blood pressure management with optimal fetal and neonatal outcomes (21). In our cohorts, we observed no major differences in *ex vivo* vascular reactivity of placental vessels between the unmedicated hypertensive group and those treated with medications during pregnancy, suggesting that *in vivo* changes in placental vascular function had a more significant impact on ex vivo vascular reactivity than exposure to maternal antihypertensive agents. These findings were unchanged following removal of pregnancies complicated by pre-eclampsia from the analysis.

In contrast, *in vivo* bisoprolol exposure was associated with a relative reduction of maximal CPA vasorelaxation to SNP of ∼26% in comparison to vessels from uncomplicated pregnancies with no medication exposure. The more pronounced impairment of vasorelaxation observed in the bisoprolol cohort may suggest that bisoprolol as a cardioselective β1-blocker alters the cell signalling pathway phenotype leading to altered eNOS expression and/or activation. It is also possible that there are chronic changes in signalling in the smooth muscle cells (e.g. impairments in NO-mediated signalling pathways) induced by *in vivo* bisoprolol exposure. Previous cohort studies support an association between *in vivo* bisoprolol exposure and changes in fetal growth (9) (53) (54) and fetal Dopplers (unpublished data from St Mary’s Hospital). However, it is very difficult to demonstrate a casual effect of bisoprolol exposure on placental vascular function as, in the same way that maternal hypertensive disease appears to influence *ex vivo* CPA vasoreactivity, there may be an underlying impact of maternal cardiac disease even though maternal blood pressure has remained normal. The maternal indications for bisoprolol use are also varied, ranging from management of maternal palpitations to significant cardiac impairment. In this small cohort, it has not been possible to disentangle the relative impact of underlying disease severity, bisoprolol duration or dose on *ex vivo* CPA function.

The regulation of the fetoplacental circulation is critical for normal fetal growth and development, as it governs oxygen, nutrient, and waste exchange between the feto-maternal compartments. Within the chorionic plate, CPAs of resistance-artery size are key determinants of fetoplacental vascular resistance and blood flow distribution, thereby influencing downstream perfusion of the villous tree and, consequently, the efficiency of oxygen and nutrient delivery to the fetus (55). Dysregulation of the fetoplacental vascular tone and/or impaired responsiveness to vasoactive stimuli can compromise fetoplacental perfusion and disrupt the exchange process within the terminal villi. This may contribute to complications such as FGR and pre-eclampsia, where placental dysfunction is central to the pathophysiology of the disease (56, 57). A multicentre retrospective cohort study showed that, compared with women not taking β-blockers, those exposed had ∼ a 1.4-fold increase in SGA and a 2.3-fold increase in FGR prevalence (9).

One limitation of this study is the concentration ranges chosen for the acute testing, which were based on known (23-29) or estimated (30, 31) fetal venous plasma concentrations. This might have overestimated physiologically relevant responses, and effects observed at pharmacological doses should be interpreted with caution.

Another limitation of this study is the heterogeneity of the participants, especially in terms of pre-delivery gestational age at which *in vivo* measurements of placental function were opportunistically recorded (and placentas obtained for *ex vivo* experiments), as well as medication dosage and timing of treatments. The length of exposure to medications varied from one week to more than one year before birth between groups, which may have had a less or more pronounced effect on CPA reactivity. For instance, the longest exposure to bisoprolol may have been more effective in downregulating endothelial paracrine signalling and/or smooth muscle cell responsivity compared to the shorter exposure to labetalol (Table S3).

Furthermore, in this study it was not possible to infer any associations between pregnancy outcomes and antihypertensive medication exposure due to significant prescriber bias and clinical presentations influencing prescribing decisions.

In conclusion, whilst cardiovascular disease in pregnancy is common the effect of antihypertensive and cardiac medications on placental vascular function remains poorly understood. Our study has confirmed the association between maternal hypertensive disorders and altered placental vascular function but has also identified differential effects of different classes of vasoactive medications in direct exposure experiments, but more importantly in vessels investigated *ex vivo* following exposure during pregnancy. Bisoprolol was associated with the most pronounced effects, justifying the need for further mechanistic studies to better inform the risk benefit discussion around the use of this medication in pregnancy.

### Perspectives

Women with hypertension and cardiovascular disease require medication to optimise maternal safety in pregnancy. Although, medications such as labetalol, nifedipine and bisoprolol are recommended to treat maternal hypertension and to minimise cardiac workload in women with cardiac disease, to date there is a lack of robust safety data to support evidence-based prescribing. We have demonstrated a significant association between maternal hypertension and a reduction in vasoreactivity of the fetoplacental vasculature which was not significantly altered by *in vivo* exposure to different antihypertensive medication classes. In addition, fetoplacental arteries isolated from normal pregnancies exposed to vasoactive drugs *ex vivo* demonstrated, that whilst CCBs are vasodilatory within the fetoplacental vasculature, adrenergic receptor agonists and antagonists have markedly different vascular effects. Fetoplacental arteries exposed to bisoprolol both *in vivo* and *ex vivo*, exhibited a vasoconstrictive phenotype. Further studies are justified to investigate the mechanisms of action of vasoactive medications in fetoplacental arteries and to understand how vasoactive mechanisms may differ from the systemic vasculature. Clinical studies are also needed to correlate *ex vivo* effects with maternal and fetal outcomes.

#### Novelty and Relevance

included as the final section of the manuscript, it should be written in a style that is understood by a general audience and is limited to about 100 words. The section must be comprised of 3 subsections.

1. **What Is New?** Fetoplacental blood arteries from women with hypertensive disorders of pregnancy have different vascular properties compared to vessels from normotensive pregnancies, but this was not altered further by chronic exposure to CCBs or labetalol. In contrast, bisoprolol was associated with an increased vasoconstrictive phenotype in arteries following acute and chronic exposure.
2. **What Is Relevant?** Different antihypertensive and cardioprotective drugs may benefit maternal health, but their pharmacological properties are not equally beneficial for fetoplacental vascular function.
3. **Clinical/Pathophysiological Implications?** Better understanding of the effects of antihypertensive and cardioprotective medications on fetal placental blood vessel function could help guide clinical practice and improve both maternal and fetal outcomes.

## Acknowledgments

We are grateful to all of the women who donated their placentas to research. We thank the staff in the Maternal and Fetal Health Research Centre for their assistance with patient recruitment, tissue and clinical data collection, particularly Jessica Morecroft.

